# Effects of pathogenic CNVs on biochemical markers: a study on the UK Biobank

**DOI:** 10.1101/723270

**Authors:** Matthew Bracher-Smith, Kimberley M Kendall, Elliott Rees, Mark Einon, Michael C O’Donovan, Michael J Owen, George Kirov

**Affiliations:** MRC Centre for Neuropsychiatric Genetics & Genomics, Institute of Psychological Medicine and Clinical Neurosciences, Cardiff University, School of Medicine, Hadyn Ellis Building, Maindy Road, Cardiff, CF24 4HQ, United Kingdom

## Abstract

**Background:** Pathogenic copy number variants (CNVs) increase risk for medical disorders, even among carriers free from neurodevelopmental disorders. The UK Biobank recruited half a million adults who provided samples for biochemical and haematology tests which have recently been released. We wanted to assess how the presence of pathogenic CNVs affects these biochemical test results.

**Methods:** We called all CNVs from the Affymetrix microarrays and selected a set of 54 CNVs implicated as pathogenic (including their reciprocal deletions/duplications) and present in five or more persons. We used linear regression analysis to establish their association with 28 biochemical and 23 haematology tests.

**Results:** We analysed 421k participants who passed our CNV quality control filters and self-reported as white British or Irish descent. There were 268 associations between CNVs and biomarkers that were significant at a false discovery rate of 0.05. Deletions at 16p11.2 had the highest number of significant associations, but several rare CNVs had higher effect sizes indicating that the lack of significance was likely due to the reduced statistical power for rarer events. The distribution of values can be visualised on our interactive website: http://kirov.psycm.cf.ac.uk/.

**Conclusions:** Carriers of many pathogenic CNVs have changes in biochemical and haematology tests, and many of those are associated with adverse health consequences. These changes did not always correlate with increases in diagnosed medical disorders in this population. Carriers should have regular blood tests in order to identify and treat adverse medical consequences early. Levels of cholesterol and related lipids were unexpectedly lower in carriers of CNVs associated with increased weight gain, most likely due to the use of statins by such people.

## INTRODUCTION

Large copy number variants (CNVs) have been shown to have significant effects on common medical disorders even among the relatively healthy individuals, with no early-onset neurodevelopmental disorders, taking part in the UK Biobank (Crawford et al, 2018). Such carriers also tended to have changes in physical traits, such as weight, height, body fat content, pulse rate and blood pressure (Owen et al, 2018).

Participants in the UK Biobank agreed to donate blood, saliva and urine samples for biochemical tests at the point of their recruitment. Biochemical and haematology tests were performed on the full set of Biobank participants and the data was recently released. We expected that there will be significant changes in the levels of many of these tests among carriers of pathogenic CNVs, in line with the increased morbidity and mortality and changes in the physical traits in this population.

## METHODS

Approval for the study was obtained from the UK Biobank under project 14421: “Identifying the spectrum of biomedical traits in adults with pathogenic copy number variants (CNVs)”.

### Participants

The UK Biobank recruited just over half a million people from the UK general population. We restricted analysis to individuals who self-reported as White British or Irish descent and whose samples passed our standard CNV QC filters (genotyping call rate < 0.96, > 30 CNVs per person, a waviness factor of < −0.03 & > 0.03 & LRR standard deviation of > 0.35).

### Choice of CNVs

We tested 54 CNVs that have been proposed to be pathogenic and were carried by at least five participants (the choice of the CNVs, their chromosomal positions and details of analysis are given in our previous publications (Crawford et al, 2018, Owen et al, 2018). Briefly, the CNVs follow the lists of Coe et al (2013) and Dittwald et al (2014).

### Tests

We analysed these CNVs for association with a set of 28 biomarkers and 23 haematology tests that were available for the majority of participants (Table 1). Details on the tests and methods used for their measurements are presented on the Biobank website: biobank.ctsu.ox.ac.uk/showcase/showcase/docs/serum_biochemistry.pdf, and biobank.ctsu.ox.ac.uk/showcase/showcase/docs/haematology.pdf. Between 401k and 319k people who passed our QC filters for CNVs had valid results for the tests (**Table 1**). Two of the available tests, Oestradiol and Rheumatoid Factor, were only performed on around 76k and 41k individuals, and were therefore excluded as they provided inadequate samples size for analysis of most rare CNVs.

**Table 1.**
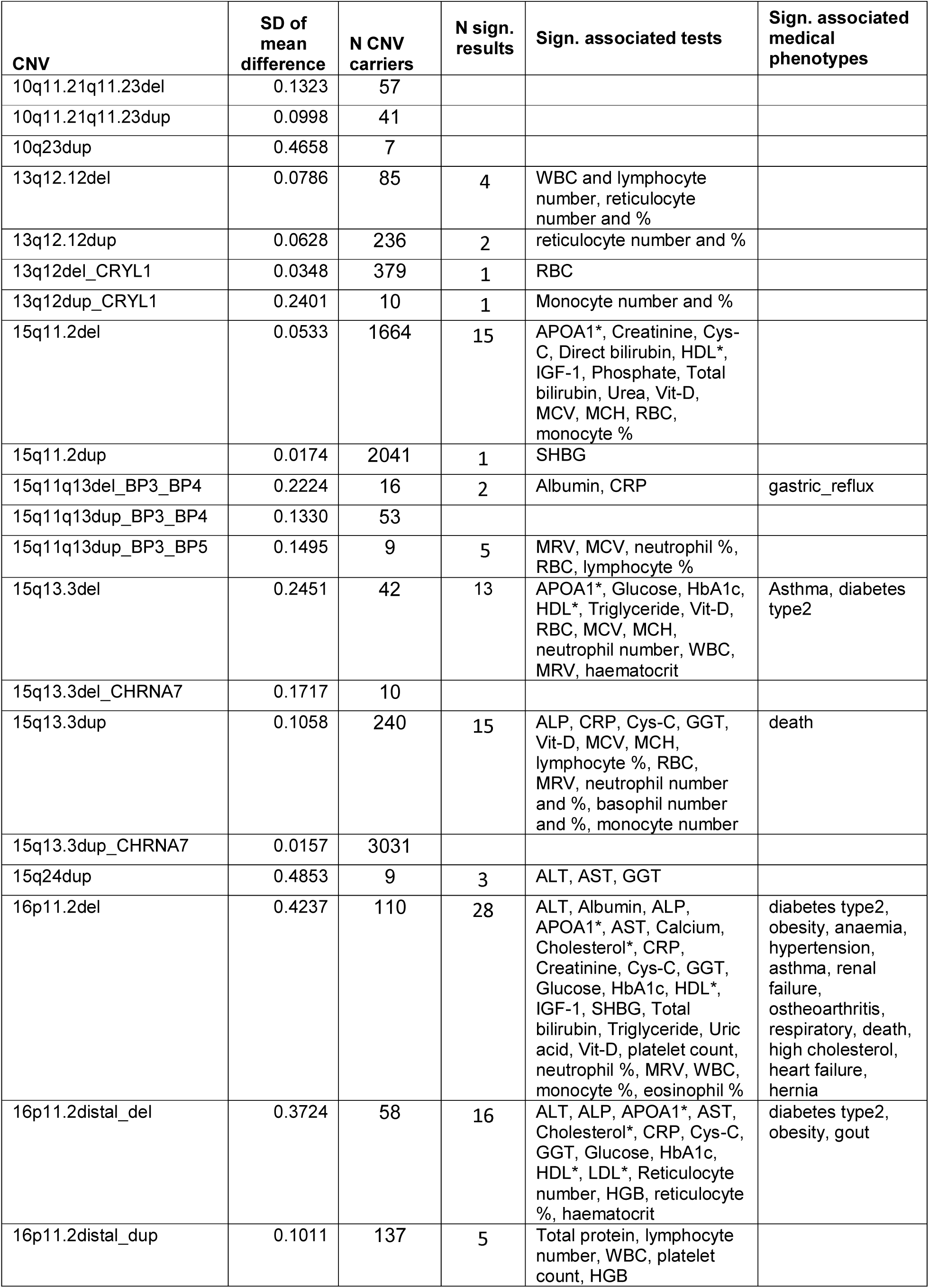
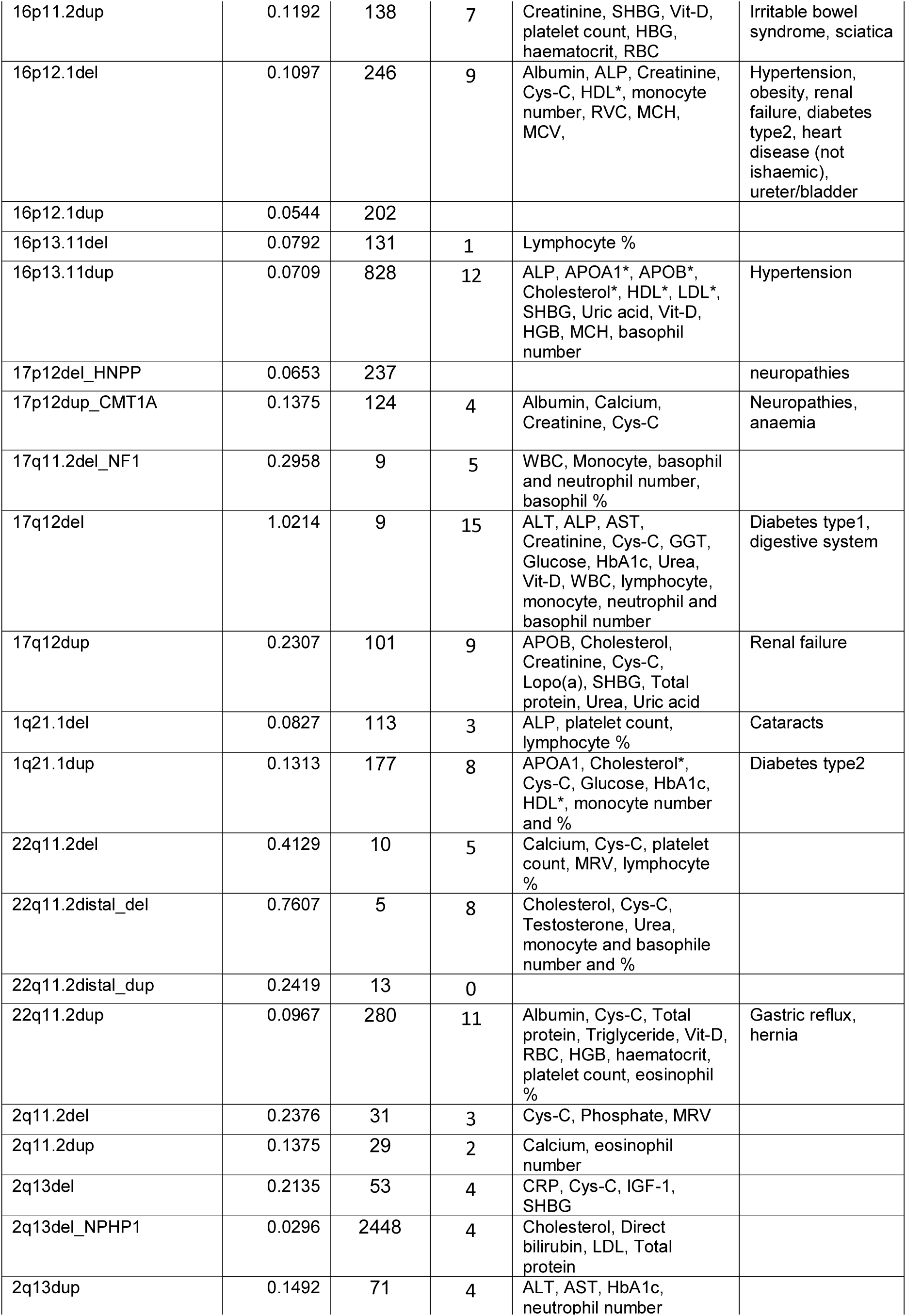
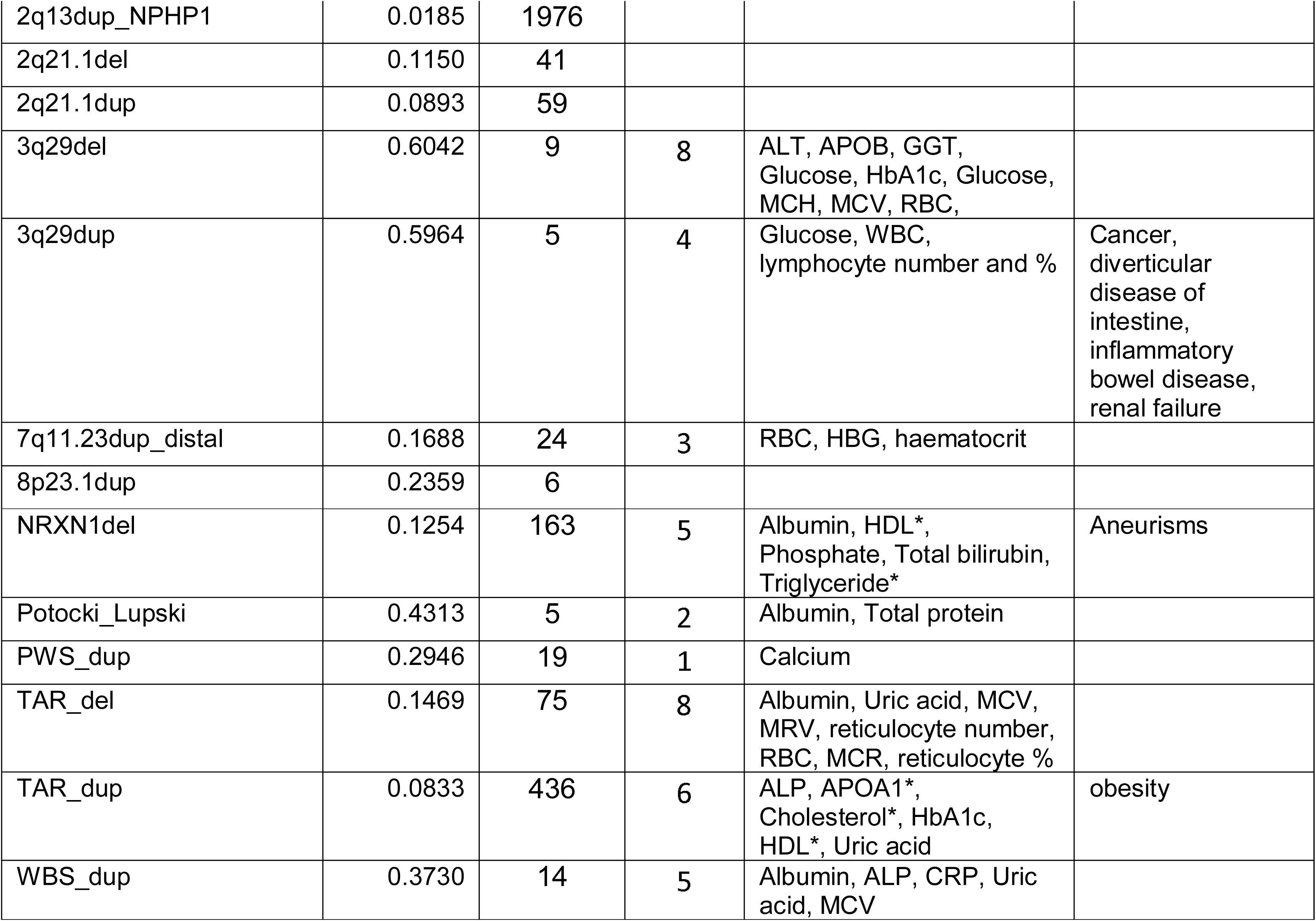
List of biochemical and haematological tests, their ranges and the number of people with valid results. The “Function” that the tests measure is listed according to the UK Biobank website.

| CNV | SD of mean difference | N CNV carriers | N sign. results | Sign. associated tests | Sign. associated medical phenotypes |
| --- | --- | --- | --- | --- | --- |
| 10q11.21q11.23del | 0.1323 | 57 |  |  |  |
| 10q11.21q11.23dup | 0.0998 | 41 |  |  |  |
| 10q23dup | 0.4658 | 7 |  |  |  |
| 13q12.12del | 0.0786 | 85 | 4 | WBC and lymphocyte number, reticulocyte number and % |  |
| 13q12.12dup | 0.0628 | 236 | 2 | reticulocyte number and % |  |
| 13q12del_CRYL1 | 0.0348 | 379 | 1 | RBC |  |
| 13q12dup_CRYL1 | 0.2401 | 10 | 1 | Monocyte number and % |  |
| 15q11.2del | 0.0533 | 1664 | 15 | APOA1*, Creatinine, Cys-C, Direct bilirubin, HDL*, IGF-1, Phosphate, Total bilirubin, Urea, Vit-D, MCV, MCH, RBC, monocyte % |  |
| 15q11.2dup | 0.0174 | 2041 | 1 | SHBG |  |
| 15q11q13del_BP3_BP4 | 0.2224 | 16 | 2 | Albumin, CRP | gastric_reflux |
| 15q11q13dup_BP3_BP4 | 0.1330 | 53 |  |  |  |
| 15q11q13dup_BP3_BP5 | 0.1495 | 9 | 5 | MRV, MCV, neutrophil %, RBC, lymphocyte % |  |
| 15q13.3del | 0.2451 | 42 | 13 | APOA1*, Glucose, HbA1c, HDL*, Triglyceride, Vit-D, RBC, MCV, MCH, neutrophil number, WBC, MRV, haematocrit | Asthma, diabetes type2 |
| 15q13.3del_CHRNA7 | 0.1717 | 10 |  |  |  |
| 15q13.3dup | 0.1058 | 240 | 15 | ALP, CRP, Cys-C, GGT, Vit-D, MCV, MCH, lymphocyte %, RBC, MRV, neutrophil number and %, basophil number and %, monocyte number | death |
| 15q13.3dup_CHRNA7 | 0.0157 | 3031 |  |  |  |
| 15q24dup | 0.4853 | 9 | 3 | ALT, AST, GGT |  |
| 16p11.2del | 0.4237 | 110 | 28 | ALT, Albumin, ALP, APOA1*, AST, Calcium, Cholesterol*, CRP, Creatinine, Cys-C, GGT, Glucose, HbA1c, HDL*, IGF-1, SHBG, Total bilirubin, Triglyceride, Uric acid, Vit-D, platelet count, neutrophil %, MRV, WBC, monocyte %, eosinophil % | diabetes type2, obesity, anaemia, hypertension, asthma, renal failure, osteoarthritis, respiratory, death, high cholesterol, heart failure, hernia |
| 16p11.2distal_del | 0.3724 | 58 | 16 | ALT, ALP, APOA1*, AST, Cholesterol*, CRP, Cys-C, GGT, Glucose, HbA1c, HDL*, LDL*, Reticulocyte number, HGB, reticulocyte %, haematocrit | diabetes type2, obesity, gout |
| 16p11.2distal_dup | 0.1011 | 137 | 5 | Total protein, lymphocyte number, WBC, platelet count, HGB |  |
| 16p11.2dup | 0.1192 | 138 | 7 | Creatinine, SHBG, Vit-D, platelet count, HBG, haematocrit, RBC | Irritable bowel syndrome, sciatica |
| 16p12.1del | 0.1097 | 246 | 9 | Albumin, ALP, Creatinine, Cys-C, HDL*, monocyte number, RVC, MCH, MCV, | Hypertension, obesity, renal failure, diabetes type2, heart disease (not ischaemic), ureter/bladder |
| 16p12.1dup | 0.0544 | 202 |  |  |  |
| 16p13.11del | 0.0792 | 131 | 1 | Lymphocyte % |  |
| 16p13.11dup | 0.0709 | 828 | 12 | ALP, APOA1*, APOB*, Cholesterol*, HDL*, LDL*, SHBG, Uric acid, Vit-D, HGB, MCH, basophil number | Hypertension |
| 17p12del_HNPP | 0.0653 | 237 |  |  | neuropathies |
| 17p12dup_CMT1A | 0.1375 | 124 | 4 | Albumin, Calcium, Creatinine, Cys-C | Neuropathies, anaemia |
| 17q11.2del_NF1 | 0.2958 | 9 | 5 | WBC, Monocyte, basophil and neutrophil number, basophil % |  |
| 17q12del | 1.0214 | 9 | 15 | ALT, ALP, AST, Creatinine, Cys-C, GGT, Glucose, HbA1c, Urea, Vit-D, WBC, lymphocyte, monocyte, neutrophil and basophil number | Diabetes type1, digestive system |
| 17q12dup | 0.2307 | 101 | 9 | APOB, Cholesterol, Creatinine, Cys-C, Lopo(a), SHBG, Total protein, Urea, Uric acid | Renal failure |
| 1q21.1del | 0.0827 | 113 | 3 | ALP, platelet count, lymphocyte % | Cataracts |
| 1q21.1dup | 0.1313 | 177 | 8 | APOA1, Cholesterol*, Cys-C, Glucose, HbA1c, HDL*, monocyte number and % | Diabetes type2 |
| 22q11.2del | 0.4129 | 10 | 5 | Calcium, Cys-C, platelet count, MRV, lymphocyte % |  |
| 22q11.2distal_del | 0.7607 | 5 | 8 | Cholesterol, Cys-C, Testosterone, Urea, monocyte and basophile number and % |  |
| 22q11.2distal_dup | 0.2419 | 13 | 0 |  |  |
| 22q11.2dup | 0.0967 | 280 | 11 | Albumin, Cys-C, Total protein, Triglyceride, Vit-D, RBC, HGB, haematocrit, platelet count, eosinophil % | Gastric reflux, hernia |
| 2q11.2del | 0.2376 | 31 | 3 | Cys-C, Phosphate, MRV |  |
| 2q11.2dup | 0.1375 | 29 | 2 | Calcium, eosinophil number |  |
| 2q13del | 0.2135 | 53 | 4 | CRP, Cys-C, IGF-1, SHBG |  |
| 2q13del_NPHP1 | 0.0296 | 2448 | 4 | Cholesterol, Direct bilirubin, LDL, Total protein |  |
| 2q13dup | 0.1492 | 71 | 4 | ALT, AST, HbA1c, neutrophil number |  |
| 2q13dup_NPHP1 | 0.0185 | 1976 |  |  |  |
| 2q21.1del | 0.1150 | 41 |  |  |  |
| 2q21.1dup | 0.0893 | 59 |  |  |  |
| 3q29del | 0.6042 | 9 | 8 | ALT, APOB, GGT, Glucose, HbA1c, Glucose, MCH, MCV, RBC, |  |
| 3q29dup | 0.5964 | 5 | 4 | Glucose, WBC, lymphocyte number and % | Cancer, diverticular disease of intestine, inflammatory bowel disease, renal failure |
| 7q11.23dup_distal | 0.1688 | 24 | 3 | RBC, HBG, haematocrit |  |
| 8p23.1dup | 0.2359 | 6 |  |  |  |
| NRXN1del | 0.1254 | 163 | 5 | Albumin, HDL*, Phosphate, Total bilirubin, Triglyceride* | Aneurisms |
| Potocki_Lupski | 0.4313 | 5 | 2 | Albumin, Total protein |  |
| PWS_dup | 0.2946 | 19 | 1 | Calcium |  |
| TAR_del | 0.1469 | 75 | 8 | Albumin, Uric acid, MCV, MRV, reticulocyte number, RBC, MCR, reticulocyte % |  |
| TAR_dup | 0.0833 | 436 | 6 | ALP, APOA1*, Cholesterol*, HbA1c, HDL*, Uric acid | obesity |
| WBS_dup | 0.3730 | 14 | 5 | Albumin, ALP, CRP, Uric acid, MCV |  |

### Statistical analysis

The tested variables followed normal distributions and did not require transformation prior to analysis. We show the results in non-normalised (original) units, to allow clinicians to relate them more easily to known tests (e.g. mmol/L, U/L). Linear regression analysis was performed with sex and age as co-variates. Other potential confounders were not included, as it is not known whether confounding factors lead to the abnormalities, or are a direct consequence of the presence of a CNV. For example, we previously showed that carrying a CNV from this list has strong adverse effects on measures of social deprivation, occupation and education (Kendall et al 2019), which are also factors that could be perceived as contributing to some biochemical tests results abnormalities. In previous work (Crawford et al, 2018) we also observed practically no effect from controlling for principal components, as should be expected for genetic variants whose frequencies are determined by selection pressure and mutation rates, rather than genetic drift (Rees et al, 2011). We used the Benjamini-Hochberg false-discovery rate (FDR) method to estimate the project-wide statistical significance (Benjamini and Hochberg 1995). A conservative false discovery rate (FDR) of 0.05 was accepted as our significance threshold (Supplementary Tables 1 and 2).

## RESULTS

The comparisons of 54 CNVs and 51 tests produced 2751 associations, of which 268 were significant at FDR=0.05 (marked in bold in Supplementary Tables 1 and 2). All results are also displayed on our institutional website (http://kirov.psycm.cf.ac.uk/) where they will be updated when required. The website allows <u>an interactive examination of the distribution of results</u>, including those for physical traits (e.g. weight and height). **Table 2** summarises the statistically significant findings. Deletions at 16p11.2 (110 carriers) produced the largest number of statistically significant associations, followed by carriers of 15q11.2 deletions, 16p11.2 distal deletions, 15q13.3 deletions and duplications and 17q12 deletions, with 15 or 16 significant associations each. **Figure 1** is as an example of the spread of test values for the top three results for 16p11.2 deletions. A substantial proportion of carriers of this CNV have abnormal results for these three tests (ranging between 18% and 44%).

**Table 2.** CNVs analysed and the associated biochemical tests and medical conditions (as presented in Crawford et al, 2018). An * denotes cholesterol and related lipid levels that are unexpectedly reduced in carriers of CNVs associated with increased BMI.

| Test | Function | N people | Mean | Min | Max | Std. Dev. | Unit |
| --- | --- | --- | --- | --- | --- | --- | --- |
| Alanine aminotransferase (ALT) | Liver | 401476 | 23.56 | 3.01 | 495.19 | 14.15 | U/L |
| Albumin | Liver | 367822 | 45.22 | 18.87 | 59.80 | 2.61 | g/L |
| Aspartate aminotransferase (AST) | Liver | 400150 | 26.22 | 3.30 | 947.20 | 10.54 | U/L |
| Direct bilirubin | Liver | 341413 | 1.83 | 1.00 | 70.06 | 0.85 | mcmol/L |
| Gamma glutamyltransferase (GGT) | Liver | 401418 | 37.45 | 5.00 | 1184.90 | 42.18 | U/L |
| Total bilirubin | Liver | 399948 | 9.13 | 1.08 | 144.52 | 4.42 | mcmol/L |
| Alkaline phosphatase (ALP) | Bone and joint | 401640 | 83.73 | 8.00 | 1416.7 | 26.53 | U/L |
| Vitamin D | Bone and joint | 383994 | 49.67 | 10.00 | 340.00 | 20.98 | nmol/L |
| Calcium | Bone and joint | 367686 | 2.38 | 1.05 | 3.57 | 0.09 | mmol/L |
| Apolipoprotein A1 (APOA1) | Cardiovascular | 365618 | 1.54 | 0.42 | 2.50 | 0.27 | g/L |
| Apolipoprotein B (APOB) | Cardiovascular | 399665 | 1.03 | 0.40 | 2.00 | 0.24 | g/L |
| Cholesterol | Cardiovascular | 401615 | 5.71 | 0.60 | 15.46 | 1.14 | mmol/L |
| C-reactive protein (CRP) | Cardiovascular | 400753 | 2.60 | 0.08 | 79.96 | 4.38 | mg/L |
| HDL cholesterol | Cardiovascular | 367652 | 1.45 | 0.22 | 4.40 | 0.38 | mmol/L |
| LDL cholesterol | Cardiovascular | 400878 | 3.57 | 0.27 | 9.80 | 0.87 | mmol/L |
| Lipoprotein (a) | Cardiovascular | 319638 | 44.11 | 3.80 | 189.00 | 49.38 | nmol/L |
| Triglycerides | Cardiovascular | 401301 | 1.76 | 0.23 | 11.28 | 1.03 | mmol/L |
| HbA1c | Diabetes | 401566 | 35.97 | 15.00 | 515.2 | 6.52 | mmol/mol |
| Glucose | Diabetes | 367387 | 5.12 | 1.01 | 36.81 | 1.21 | mmol/L |
| Creatinine | Renal | 401421 | 72.29 | 10.80 | 1499.3 | 17.95 | mcmol/L |
| Cystatin C | Renal | 401577 | 0.91 | 0.30 | 7.49 | 0.17 | mg/L |
| Phosphate | Renal | 367125 | 1.16 | 0.37 | 4.70 | 0.16 | mmol/L |
| Total protein | Renal | 367406 | 72.36 | 36.27 | 117.36 | 4.04 | g/L |
| Urea | Renal | 401346 | 5.43 | 0.81 | 41.83 | 1.39 | mmol/L |
| Uric acid (UA) | Renal | 401135 | 309.39 | 89.10 | 884.30 | 80.41 | mcmol/L |
| Insulin-like Growth Factor-1 (IGF-1) | Cancer | 399465 | 21.37 | 1.45 | 126.77 | 5.66 | nmol/L |
| Sex Hormone Binding Globulin (SHBG) | Cancer | 364235 | 51.90 | 0.39 | 241.92 | 27.72 | nmol/L |
| Testosterone | Cancer | 363579 | 6.57 | 0.35 | 54.34 | 6.05 | nmol/L |
| Basophill number | haematology | 407888 | 0.00 | 2.60 | 0.03 | 0.05 | x 10<sup>9</sup> cells/L |
| Basophill percent | haematology | 407894 | 0.00 | 33.80 | 0.57 | 0.61 | percent |
| Eosinophill number | haematology | 407888 | 0.00 | 9.60 | 0.17 | 0.14 | x 10<sup>9</sup> cells/L |
| Eosinophill percent | haematology | 407894 | 0.00 | 100.00 | 2.56 | 1.85 | percent |
| Haematocrit | haematology | 408607 | 0.05 | 72.48 | 41.12 | 3.52 | percent |
| Haemoglobin concentration (HGB) | haematology | 408607 | 0.11 | 22.27 | 14.20 | 1.23 | g/dl |
| Lymphocyte number | haematology | 407888 | 0.00 | 177.00 | 1.95 | 1.16 | x 10<sup>9</sup> cells/L |
| Lymphocyte percent | haematology | 407894 | 0.00 | 98.70 | 28.65 | 7.36 | percent |
| Mean corpuscular volume (MCV) | haematology | 408605 | 53.52 | 160.30 | 91.33 | 4.39 | fL |
| Mean corpuscular haemoglobin (MCH) | haematology | 408604 | 0.00 | 95.67 | 31.55 | 1.83 | g/dL |
| Mean corpuscular haemoglobin concentration (MCHC) | haematology | 408600 | 16.10 | 97.30 | 34.54 | 1.06 | fL |
| Mean reticulocyte volume (MRV) | haematology | 402048 | 46.00 | 249.45 | 105.84 | 7.77 | fL |
| Monocyte number | haematology | 407888 | 0.00 | 34.26 | 0.48 | 0.22 | x 10<sup>9</sup> cells/L |
| Monocyte percent | haematology | 407894 | 0.00 | 96.90 | 7.09 | 2.70 | percent |
| Neutrophill number | haematology | 407888 | 0.00 | 52.02 | 4.25 | 1.41 | x 10<sup>9</sup> cells/L |
| Neutrophill percent | haematology | 407894 | 0.00 | 97.70 | 61.13 | 8.40 | percent |
| Nucleated red blood cell number | haematology | 407878 | 0.00 | 6.90 | 0.00 | 0.03 | x 10<sup>9</sup> cells/L |
| Nucleated red blood cell percent | haematology | 407874 | 0.00 | 48.86 | 0.03 | 0.40 | percent |
| Platelet count | haematology | 408605 | 0.30 | 1821.0 | 253.44 | 59.89 | x 10<sup>9</sup> cells/L |
| Red blood cell count | haematology | 408607 | 0.01 | 7.91 | 4.51 | 0.41 | x 10<sup>12</sup> cells/L |
| Reticulocyte number | haematology | 402047 | 0.00 | 2.39 | 0.06 | 0.04 | x 10<sup>12</sup> cells/L |
| Reticulocyte percent | haematology | 402047 | 0.00 | 90.91 | 1.35 | 0.89 | percent |
| White blood cell count (WBC) | haematology | 408602 | 0.00 | 189.50 | 6.90 | 2.06 | x 10<sup>9</sup> cells/L |

**Figure 1.**
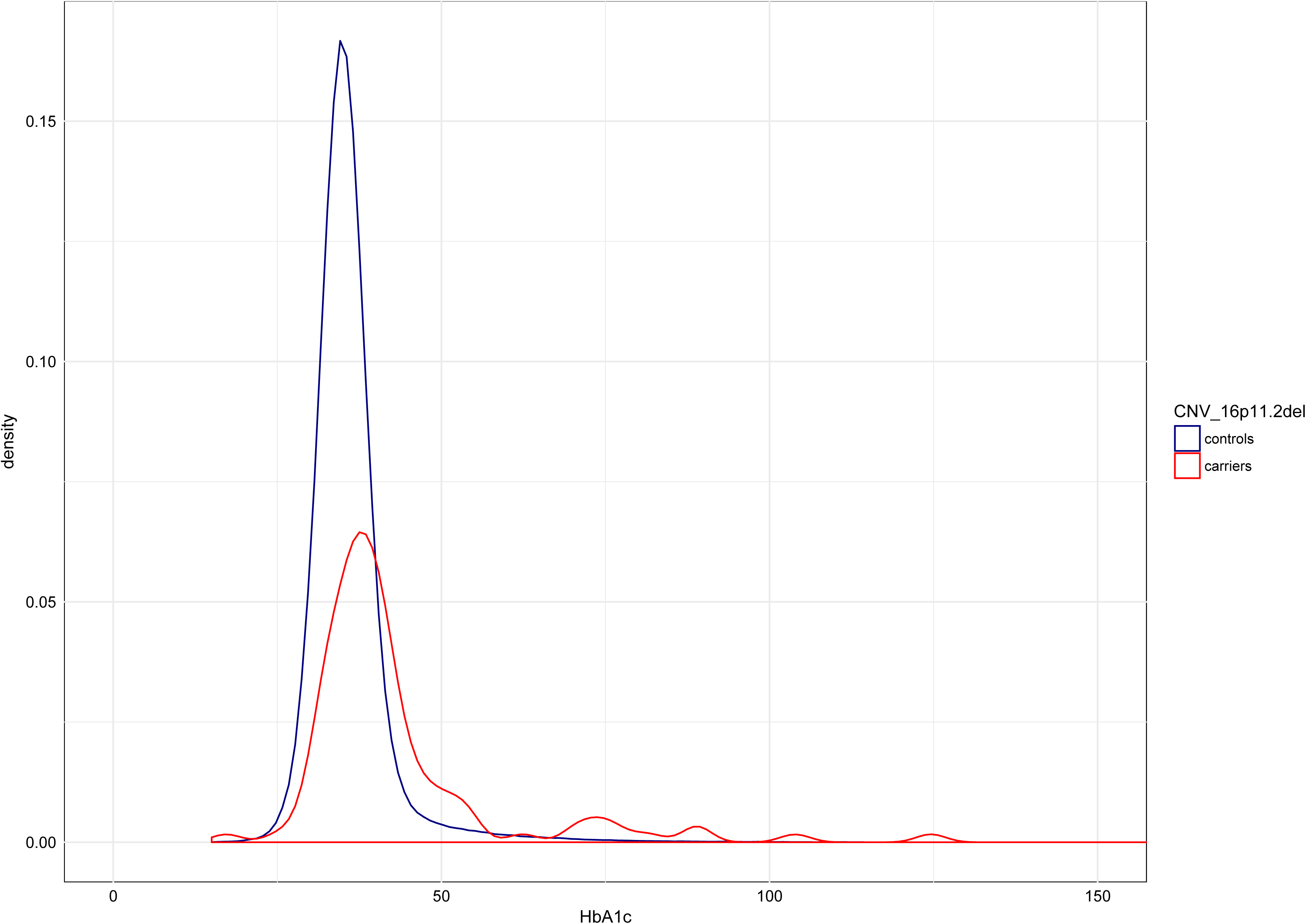
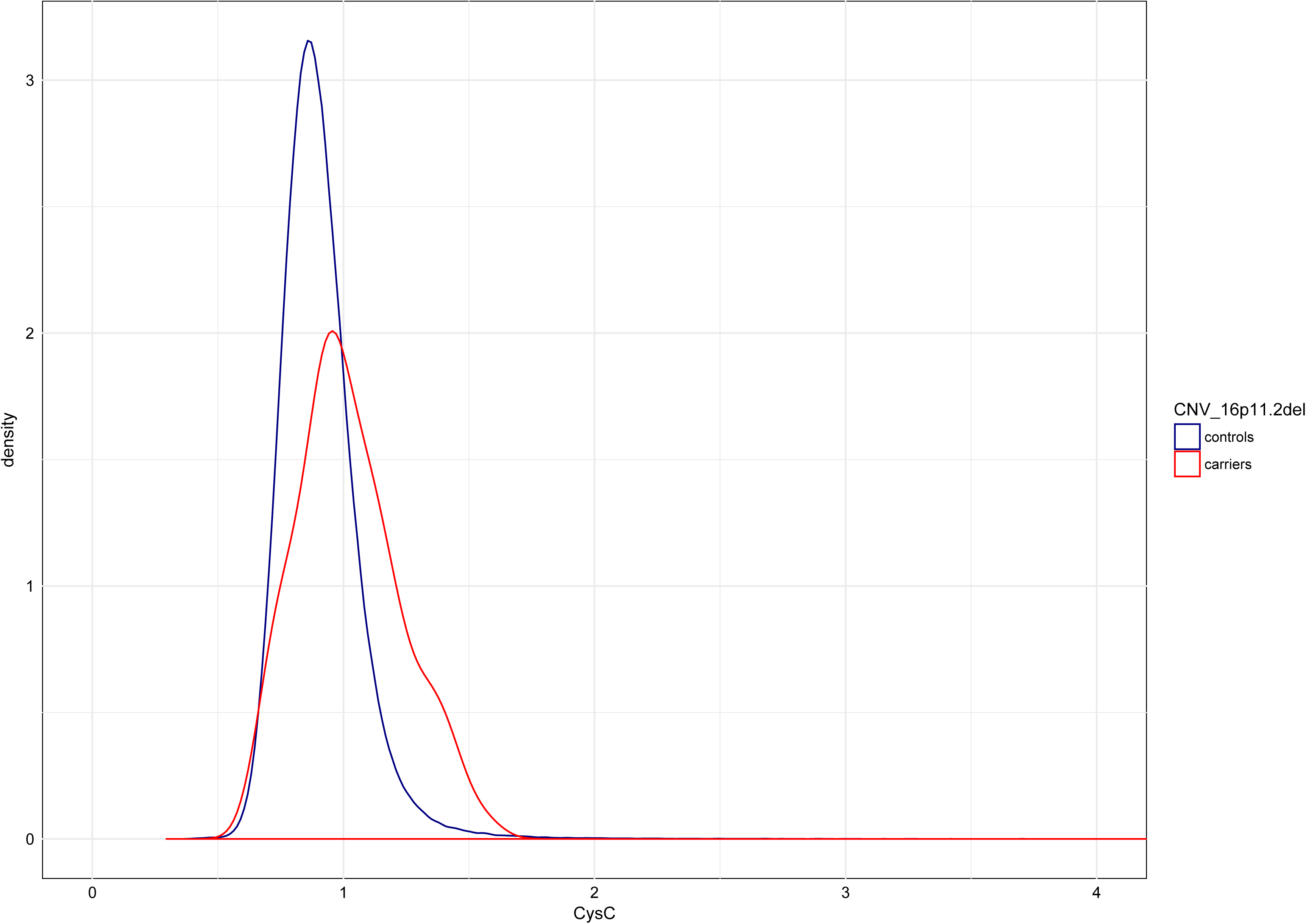
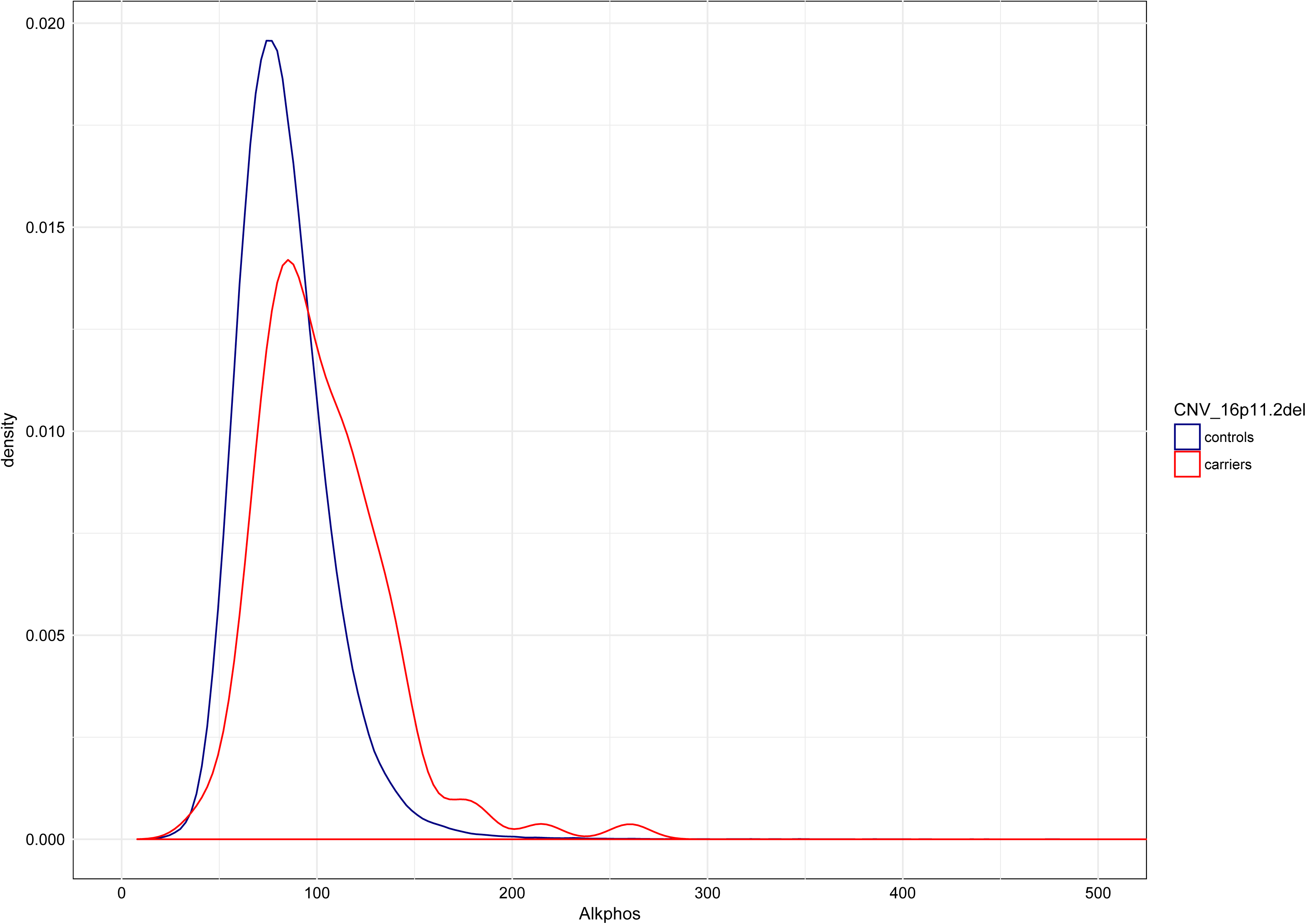
Spread of the values of the top three findings for 16p11.2 deletions and controls: a) HbA1c. Normal levels of HbA1c are ≤40mmol/mol, leaving 43% of CNV carriers with increased levels. b) Cystatin-C. Normal levels for Cystatin C are ≤1.02 mg/L for people over the age of 50, leaving 44% with abnormal levels. c) Alkaline phosphatase. Normal levels for Alkaline Phosphatase are <130 U/L for adults, leaving 18% of carriers with abnormal levels.

Several rarer CNVs had even larger average differences compared to controls (in terms of standard deviations), which did not always reach significance, very likely due to the smaller sample sizes. The most pathogenic ones appear to be some of the rarest CNVs, with 5-10 carriers each: 17q12 deletions (Renal Cysts and Diabetes syndrome), 17p11.2 duplications (Potocki-Lupski syndrome), 3q29 deletions and duplications, 22q11.2 classic and distal deletions, 15q24 duplications and 10q23 duplications.

Some of the well-established medical consequences of CNVs were associated with corresponding abnormalities in biochemical tests. For example 17q12 deletions (Renal Cysts and Diabetes syndrome) were associated with increased levels of creatinine, urea, Cystatin-C and glucose; 22q11.2 deletion carriers exhibited reduced calcium levels; 16p11.2 deletion carriers had the expected multiple abnormalities associated with cardiometabolic disorders shown for this condition; and 16p12.1 deletion carriers, who have high rates of renal failure and heart disease, have abnormalities in creatinine and Cystatin-C (**Table 2**). Other abnormalities were more subtle and would have escaped detection, if they had not been analysed in such a large dataset. Most notable are the many changes detected among carriers of 15q11.2 deletions, a CNV with neurodevelopmental phenotypes, but no confirmed medical co-morbidities. The changes were subtle (around 0.1 SD for the significant results and 0.05 SD overall) and therefore unlikely to result in significant increases of overt medical disorders. As an example, 23.7% of carriers had increased levels of Cystatin-C, compared to 18% of non-carriers. However, the abnormalities might manifest with problems in later life and reduce life expectancy if left untreated.

Most but not all significant changes suggest deterioration of function, as follows: **Liver**: Nearly all significant changes affecting liver function test (except bilirubin levels) were in the direction of higher levels, indicating liver abnormalities. **Bone**: All significant results for the three tests connected with bone function were adversely changed - reduced Vitamin D and Calcium, raised ALP. **Diabetes**: All significant changes were in the direction of raised glucose and HbA1c. **Renal**: All significant Cystatin-C results (in 14 CNVs) were increased, as were nearly all significant urea and uric acid levels, suggesting reduced renal function in several CNVs. There were exceptions: 17p12 duplications (causing Charcot-Marie-Tooth disease type 1A) and 16p11.2 deletion carriers had reduced creatinine levels. **Cancer markers**: There were very few significant results, with mixed effects. It is difficult to interpret these as a group, as they are relevant for specific cancers. **Haematology**: Apart from general trends for increases in neutrophil and lymphocyte counts, there were no clear patterns. Several CNVs showed changes predominantly in haematological test results (as opposed to biochemistry results): 13q12.12 and 15q13.3 deletions and duplications, 15q11q13_BP3_BP5 duplications, 16p11.2 distal duplications, 17q11.2 (NF1) deletions, 7q11.23 distal duplications and TAR deletions. **Cardiovascular**: Results from several tests were deemed uninformative in this population, (most likely due to statins intake, as explained in the Discussion).

Homozygous deletions at 2q13del locus affect the gene NPHP1 and are known to cause the kidney disorder *juvenile nephronophthisis*. Three homozygous individuals were excluded from analysis, as we have reported previously that all three had renal failure (Crawford et al, 2018). Heterozygous individuals have subtle phenotypes, with changes in cholesterol and total protein (reductions) and increases in direct bilirubin. Creatinine was also increased but the evidence for association was weak after correction for multiple testing, with a change of less than 1 mcmol/L.

Mirror-image phenotypes, as reported for physical traits in our previous work (Owen et al, 2018) were rare, almost entirely confined to 16p11.2 (creatinine, platelet count and SHBG) and for 15q13.3 (MCH, MCV and MRV).

## DISCUSSION

Most pathogenic CNVs tested in this study are known to increase risk for neurodevelopmental disorders and common medical phenotypes. As such it is not surprising that they are also associated with abnormal biochemical tests. About 10% of all possible CNV/test associations were significant at a conservative FDR = 0.05, suggesting multiple effects. While most are expected, e.g. high creatinine and Cystatin-C for CNVs associated with renal failure, others suggest more subtle effects that could affect medical outcomes and all-cause mortality, some of which have so far not been demonstrated. Indeed, we previously showed that carriers of 11 of these CNVs have statistically significant increase in mortality, which was not fully explained by the statistically significant associated medical conditions (Crawford et al, 2018).

One set of associations runs against the expected directions and deserves a further discussion: Cholesterol and associated biochemical tests (LDL, HDL, ApoA1 and ApoB) were *reduced* (rather than increased) among carriers of CNVs that are associated with obesity or ischaemic heart disease, such as 16p12.1 deletions, 16p11.2 deletions and 16p11.2 distal deletions (Bochukova et al, 2010, Crawford et al, 2018). We suspect that this is due to a large proportion of such patients taking statins, precisely because they are overweight or have cardiovascular problems. This possibility is confirmed by an extreme example: people diagnosed with “high cholesterol” in the sample as a whole, also showed significantly reduced levels of these tests, a seemingly counter-intuitive finding. Levels of cholesterol were, on average, 0.66 mmol/L lower among the 70,052 people who had this diagnosis, a situation only likely if statins were taken after the diagnosis was made (**Table 3**). This class of medicines is highly effective in lowering cholesterol and lipid concentrations (Cholesterol treatment trialists, 2019). At the point of recruitment the participants listed the medications they were taking, so we tested whether reductions in cholesterol were present only among those on statins. This could not fully explain the observations. Even CNV carriers who had not declared statins intake had lower cholesterol levels than non-carriers who were not taking statins. In a linear regression analysis, being on statins was associated with a 1.61 mmol/L lower cholesterol for the population as a whole, while a diagnosis of “high cholesterol” was associated with only 0.4 mmol/L higher cholesterol. Similar analysis on 16p11.2 deletion carriers showed that being on statins was associated with a 1.34 mmol/L lower cholesterol, while carrying the deletion was associated with 0.19 mmol/L lower cholesterol. We suggest that the only way to reconcile these findings is that a proportion of people (CNV carriers and non-carriers) had either taken statins at an earlier time, or had not declared them. CNV carriers who are obese or have cardiovascular incidents could be more likely to have been prescribed statins. A separate analysis of all biochemical results with statins intake as a co-variate are presented in **Supplementary Table 2**. Researchers should be aware of potential bias in the UK Biobank when interpreting findings on cholesterol and other lipids, a problem which could be reduced once the primary care (GP) data become available.

**Table 3.** Cholesterol levels in cases and controls, according to statins intake, diagnosis of “high cholesterol” and presence of a 16p11.2 deletion, an example of a CNV highly associated with obesity and cardiometabolic disorders.

| Group | On statins | Cholesterol mean (SD) | N participants |
| --- | --- | --- | --- |
| People with no diagnosis of “High cholesterol” | No | 5.89 (1.05) | 316856 |
|  | Yes | 4.39 (0.89) | 14707 |
| People with a diagnosis of “High cholesterol” | No | 6.38 (1.23) | 19153 |
|  | Yes | 4.74 (0.93) | 50899 |
| CNV non-carriers | No | 5.92 (1.07) | 323170 |
|  | Yes | 4.66 (0.94) | 63000 |
| 16p11.2del carriers | No | 5.58 (1.31) | 78 |
|  | Yes | 4.45 (1.14) | 25 |

## CONCLUSIONS

Our findings of changes in biochemical tests extend our previous reports on this population, showing adverse changes in basic physical characteristics in CNV carriers (Owen et al, 2018) and higher rates of common medical disorders (Crawford et al, 2018). This strengthens the case for general monitoring of such carriers and treatment and prevention of biochemical abnormalities, such as with statins or antidiabetic drugs. As an example, 43% of carriers of 16p11.2 deletions have levels of HbA1c above the normal range of 40 mmol/mol, compared to 12.7% of CNV non-carriers (**Figure 1**). The findings of this study cannot yet be used for analysis of cholesterol and related lipids, due to the high rates of statins intake in this population. Clinicians involved in the care of CNV carriers will be able to examine the results in more detail, using our interactive website which shows the distribution of values between carriers and non-carriers, separately for men and women: http://kirov.psycm.cf.ac.uk/.

## Acknowledgements

This research has been conducted using the UK Biobank Resource under Application Number 14421.

## Conflict of interest statement

The work at Cardiff University was funded by the Medical Research Council (MRC) Centre Grant (MR/L010305/1) and Program Grant (G0800509).

## Notes

#### Summary of Updates

An interactive website is now added, to allow readers to visualise the distribution of values of biochemical markers and physical traits among carriers and non-carriers of CNVs. It was developed by the first author Matthew Bracher-Smith

https://kirov.psycm.cf.ac.uk/

